# Cytoskeletal mechanisms of axonal contractility

**DOI:** 10.1101/227777

**Authors:** S. P. Mutalik, J. Joseph, P.A. Pullarkat, A. Ghose

## Abstract

Mechanotransduction is likely to be an important mechanism of signalling in thin, elongated cells like neurons. Maintenance of pre-stress or rest tension may facilitate mechanotransduction in these cells. In recent years, functional roles for mechanical tension in neuronal development and physiology are beginning to emerge but the cellular mechanisms regulating neurite tension remain poorly understood. Active contraction of neurites is a potential mechanism of tension regulation. In this study, we have explored cytoskeletal mechanisms mediating active contractility of neuronal axons. We have developed a simple assay where we evaluate contraction of curved axons upon trypsin-mediated detachment. We show that curved axons undergo contraction and straighten upon de-adhesion. Axonal straightening was found to be actively driven by actomyosin contractility, while microtubules may subserve a secondary role. We find that while axons show a monotonous decrease in length upon contraction, subcellularly, the cytoskeleton shows a heterogeneous contractile response. Further, using an assay for spontaneous development of tension without trypsin-induced de-adhesion, we show that axons are intrinsically contractile. These experiments, using novel experimental approaches, implicate the axonal cytoskeleton in tension homeostasis. Our data suggest that while globally the axon behaves as a mechanical continuum, locally the cytoskeleton is remodelled heterogeneously.

## INTRODUCTION

Neurites are axi-symmetric structures with extreme aspect ratio and long lengths. Developing neurites experience mechanical stretch via growth cone mediated towing (1) and later, after the formation of connections, from tissue expansion associated with animal growth. Mechanical tension along neurites has been demonstrated to affect various neuronal processes including growth (2), retraction (3), synaptic vesicle dynamics (4, 5), synaptic transmission (6), excitability (7) and network formation (8, 9). Though the focus of the axon mechanics field has been on axonal stretch as a feedback mechanism for various biological processes, not much is known about the mechanisms maintaining and regulating neurite tension.

The role of axonal cytoskeleton, its organization and dynamics in the regulation of tension, is beginning to emerge. The axonal shaft is filled with heavily cross-linked polarized bundles of microtubules with plus-end oriented towards growth cone (10). F-actin is organized by crosslinkers and myosins into a thin cortical layer underneath the plasma membrane (11). In mature neurites, actin and spectrin form a periodic, membrane-associated skeleton rings (12, 13), and together with microtubules are known to protect neurites from mechanical stress (14). Other organisations of F-actin like, actin trails, patches and dynamic actin hotspots have also been reported in axons (15). These recent observations and others show that the axonal cytoskeleton is highly dynamic and it is likely that remodeling of the cytoskeleton and membrane-cytoskeleton complexes will be central to the regulation of axonal tension.

Several studies have demonstrated that neurites, including axons, actively maintain a rest tension (2, 16, 17). Neuronal rest tension has largely been studied using microneedle-based manipulations (2, 18) and ablation studies (16). In microneedle-based axon slackening experiments, the recovery tension often exceeded the initial values suggesting active generation of tension in neurons (2). In axotomy-induced retraction based studies, axonal shortening was found to be ATP-dependent (16). Later, laser ablation induced axonal retraction was demonstrated to be dependent on actomyosin contractility (19). In PC12 neurites, retraction studies revealed that actin was involved in generating tension while microtubules had a resisting function (20). A similar balancing function between actomyosin contractility and microtubule resistance has been recently identified in contracting *Drosophila* neurons (21). Active contraction as a potential mechanism to regulate axonal rest tension has been suggested from studies on PC12 neurites and modelled by invoking active involvement of molecular motors (18).

Most of the above mentioned studies have employed acute and localized perturbations, which may invoke local responses to these perturbations or damage. For example, axotomy is known to locally elevate calcium (22) and, in turn, may induce activation of myosins (19) and calpain proteases (23). A study attempting to circumvent these issues employed flow stretching of PC12 neurites and reported oscillations in the strain rate suggestive of active behavior (24). In the current study, we have developed simple assays that are globally acting and non-intrusive compared to microneedle or ablation strategies. We use these methods to investigate the origins of axonal contractility in vertebrate sensory neurons. Our results suggest that axons are intrinsically contractile and axonal contraction is driven primarily by actomyosin activity.

## MATERIALS AND METHODS

### Dissection and cultures

Fertilized, white Leghorn chicken eggs were obtained from Venkateshwara Hatchery Private Limited, Pune, India. All procedures were approved by the Institutional Animal Ethics Committee, IISER Pune, India. Nine or ten-day old chick embryos were used to isolate dorsal root ganglia (DRG). Dissections were carried out in sterile PBS (137mM NaCl, 2.7mM KCl,10mM Na_2_HPO_4_,1.8mM KH_2_PO_4_) under a dissecting microscope inside a horizontal laminar flow hood. Twelve to fourteen DRGs were removed from both sides of the spinal cord and collected in L15 medium (Gibco). The tissue was centrifuged at 3000 rpm for 3 minutes before the medium was removed and 1x trypsin (Lonza) was added. Trypsinization was undertaken for 20 minutes at 37 °C. Following trypsinization, the tissue was centrifuged at 3000 rpm for 3 - 5 minutes before removal of trypsin and washed with L15 medium. The dissociated neurons were plated on poly-L-lysine-coated (1mg/ml PLL; Sigma) 35 mm cover glass bottomed petri dishes in 1.5 ml of serum-free media (L15 containing 6 mg/ml glucose (Sigma), 1x glutamine (Gibco), 1x B27 (Invitrogen), 20 ng/ml NGF (Gibco) and 1x Penstrep (Gibco)). The neuronal culture was incubated for 48 hours at 37 °C before de-adhesion experiments. Cells were grown in serum-free media until half an hour before de-adhesion experiments in order to increase the ocurrance of curved axons during the initial growth phase (see next section for details).

For micro-contact printing experiments, the cover glasses were patterned (as described below) before plating the neurons.These experiments were performed in media containing 10% fetal bovine serum (Gibco).

### Trypsin de-adhesion, imaging and analysis

In initial experiments, we grew DRG neurons in the presence of serum prior to trypsin deadhesion. However, subsequently, we took advantage of the fact that neurons cultured without serum have a much greater frequency of neurites showing curved trajectories (25) and thus improved the throughput of our experiments. Neurons were grown for 48 hours without serum followed by re-feeding 10% FBS 30 minutes or 2 hours prior to trypsinisation. In all three conditions (viz., with serum or serum-free followed by 30 minutes or 2 hours serum re-feeding) straightening response was seen in response to axonal de-adhesion (Figure S1).The contractility of 30 minutes and 2 hours serum re-fed axons were comparable. Thus, all inhibitor-based perturbation experiments were conducted following 30 minutes of serum treatment.

To perform trypsin de-adhesion experiments, 10% fetal bovine serum (Gibco) was added to the neurons grown in serum-free media and incubated for a further 30 minutes at 37 °C. Immediately prior to trypsin-induced de-adhesion, serum containing media was removed, the cultures were washed thrice in prewarmed serum-free L15 and replaced with 1 ml of serum-free L15. Isolated curved axons were identified and trypsin (Sigma) was added to a final concentration of 5x. Upon trypsin treatment, axons de-adhere and show a straightening response. The axonal de-adhesion precedes the detachment of soma and growth cones from the substrate. Neurons were imaged at 37 °C usingDifferential Interference Contrast (DIC) microscopy using a 40x oil objective on an Olympus IX81 system equipped with a Hamamatsu ORCA-R2 CCD camera. Images were recorded at 1 frame per second acquisition rate using the Xcellence RT (Olympus) software. Typically, straightening starts after 10-20 seconds of trypsin addition and imaging was continued till there is no further length change or the axonal ends (soma or growth cone) detach. Images were exported to ImageJ and lengths were measured using the Segmented Line tool at every 10-second interval.

The following inclusion criteria were used to select axons for evaluation of axonal contractility.

1. Axons showing significant curvature, presumably due to attachments along their lengths, were chosen for axon straightening experiments (Figure 1 A). Axonal segments having obvious branches were not considered as the branches may hinder contraction.
2. If the growth cone retracted concomitantly with a reduction in axonal curvature then such data were included in our analysis. In these cases the assumption is that the axonal shortening, evident from the reduction of curvature, is causing the weakly attached growth cone to be pulled backwards (Figure 1B).
3. If the growth cone or soma detached and retracted before the reduction in axonal curvature then these were not included in the analysis.

In these experiments, the axon lengths ranged from 70 μm to 180 μm and 5 - 35% decrease in length was observed upon de-adhesion.

### Neurons grown on patterened substrates

Patterned substrates were generated using micro-contact printing. Silicon masters with 20 μm diameter depressions spaced 50 μm or 70 μm apart were procured from Bonda Technology Pte. Ltd., Singapore. PDMS (Dow Corning) stamps were made from the master by using a previously published protocol (26). Sterile PDMS stamps were washed with isopropanol and dried in the laminar flow hood. Laminin (20 μg/ml), fibronectin (100 μg/ml) and rhodamine-labeled BSA (10 μg/ml) were mixed and used for “inking” the PDMS stamp. The protein mixture was applied onto the stamp and incubated for 5 minutes at room temperature. After removal of excess protein solution, the stamps were used to pattern cover glass bottomed 35 mm petri dishes. PBS was added immediately after printing to avoid drying of the patterns until neurons were plated.

For studies involving axons straightening on patterned substrates, axons were imaged after 8 hours or overnight incubation in serum containing medium. Neurons were imaged at 37 °C in DIC mode using 10x or 40x objective at 15-second intervals. Images were exported to Image J and axonal lengths measured using the segmented line tool.

### Drug treatments

Neurons were treated with 30 μM Blebbistatin (- enantiomer; Sigma) for 1 hour prior to the straightening experiments. For actin de-polymerization and microtubule depolymerization experiments, 0.6 μM Latrunculin A (Sigma) or 16 μM / 33 μM Nocodazole (Sigma) were added to the cultures 15 minutes prior to trypsin treatment. All drugs were dissolved in DMSO. Control experiments were undertaken to test the effect of similar volumes of DMSO on straightening kinetics.

Efficacy of Latrunculin A and Nocodazole treatment was confirmed using phalloidin staining and alpha tubulin immunofluorescence, respectively (see Supplementary Materials and Methods for details).

### Labeling of mitochondria

Neurons were cultured for 48 hours followed by serum induction as described above. After 30 minutes of serum induction, cells were incubated with 50 nM MitoTracker Green FM (Thermo Scientific) for 2 minutes followed by 2 washes using L15 (Gibco) and minimum 30 minutes incubation in L15 before imaging. Images were acquired at every one second interval in GFP channel using a 40x oil objective on an Olympus IX81 system equipped with a Hamamatsu ORCA-R2 CCD camera.

### Mitochondria tracking and analysis

We developed a MATLAB-based code to track labeled mitochondria. Mitochondria were tracked every 10 seconds as this was optimal for differentiating docked and mobile mitochondria. Simple Neurite Tracer plugin (http://imagej.net/Simple_Neurite_Tracer) was implemented in Fiji (https://imagej.net/Fiji) to trace the axon and obtain the coordinates along its length at each time point. These images, with their corresponding coordinate files were saved as .jpg and .swc files, respectively. Images and their corresponding .swc files were imported to MATLAB. Coordinates from the .swc files were used to obtain the normal to the path forming the neurite trajectory at each point and the vector of pixels along this normal was determined. The location of the intensity maxima along this vector of pixels at each point was used as the coordinate of the neurite at that point. The vectors of pixels along the normals were also stored for making kymograph plots. The intensity values per pixel along the neurite, using the coordinates obtained by the procedure described above, were used to identify the locations of the mitochondria. To eliminate detection of small spurious local peaks as mitochondria a smoothing filter was used. Intensity series from all images of a neurite were resampled to have the same length as the longest (the first image). These interpolated series were used to reliably identify peaks corresponding to mitochondria that were consistently present throughout the experiment (Figure S6). The coordinates of the detected mitochondria were scaled back to get their coordinates in the original scale. These position data were used to calculate local strains.

### Definitions of parameters

We characterized the length minimization response by evaluating the evolution of the strain in the axonal segment (Figure 1A,B). 
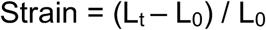

For quantitative comparison of data under various cytoskeleton pertubing drug treatments we have used contraction factor (C_r_) as a time-independent measure of the extent of axonal shortening (21) (Figure S2), 
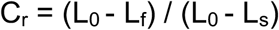

L_f_: = saturation length of the axon after de-adhesion, measured form the last frame of recording.
L_0_: = length of the axon at the time of trypsinization (t = 0).
L_s_: = straight line distance between end points of axonal segment at time 0.
L_t_: = length of the axon at time t

If there is no length change then C_r_ will be 0, C_r_ = 1 indicates straight end-to-end geometry, and values between zero and one indicates partial straightening. And if length reduction is accompanied by tip retraction then C_r_ could exceed 1.

### Statistics

Curve fitting and plotting of strain vs time plots were done in Matlab (2014) using the cftool. Average strain plots were plotted in Excel (2007 or 2016). Unpaired t-test was used to compare contraction factors using the GraphPad Prism 5 software.

## RESULTS AND DISCUSSION

### Axons show strain relaxation and straightening upon trypsin-mediated detachment

In order to evaluate the axonal strain without acute mechanical perturbations or damage, we developed an assay involving straightening of curved neurites in culture. In this assay, primary chick DRG neurons are cultured on poly-L-lysine coated glass substrates and axonal strain is evaluated following trypsin-induced axonal de-adhesion. Upon addition of 5x trypsin, axons detach faster from the substrate than the growth cone or the neuronal soma. Presumably, the relatively smaller contact surface of the axon supports fewer attachments compared to the growth cone or cell body. We find that in response to trypsin-mediated axonal de-adhesion, straightening is evident in the axonal segment and progressive axonal shortening results in length minimization (Figure 1A-C; Movie S1). Typically, the reduction in curvature is apparent within 10-20 seconds of trypsin addition and it takes 1 - 5 minutes for the axonal segment to reach the minimum length.

In order to evaluate the length shortening response, we used strain as a parameter (Strain = (L_t_ − L_0_) / L_0_ where L_0_, length of the axon at the time of trypsinization (t = 0) and L_t_, length of the axon at time t). Strain is 0 at the time of trypsin addition and the negative strain reflects the length shortening response. Interestingly, neurons show heterogenous behavior. Some neurons showed a linear change in strain with a constant rate (Figure 2A) while the other neurons displayed an exponential strain relaxation response (Figure 2B). In the latter case, the strain saturates at different values. These different trends could be due to underestimation of the data for neurons showing a linear response because of de-adhesion of the ends as an axon straightens or differences in extent of length change. It is also possible that these differences arise from heterogeneity in intrinsic contractility within individual neurons. Heterogeneity in recovery of rest tension following slackening has been previously documented in chick DRGs (2). While some neurons recovered to tension values higher than the initial, others failed to reach the initial value. Further, repeated slackening of the same neurons suggested a tendency to behave similarly (2). These observations were independent of initial length or initial tension and reinforce the possibility of intrinsic differences in contractility between neurons.

**Figure 1.**
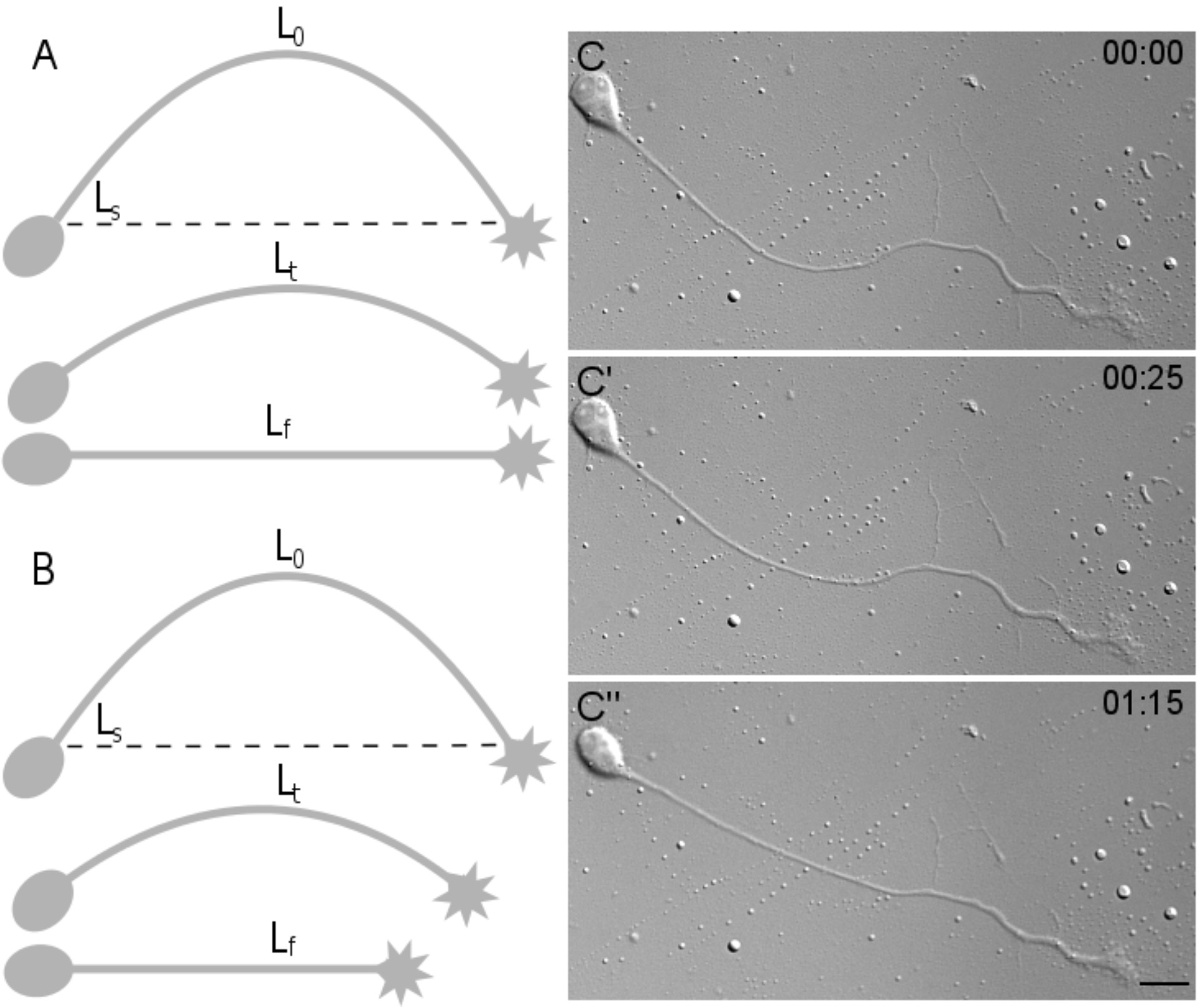
Trypsin induced axonal de-adhesion and straightening. (A) Schematic representation of axonal contraction upon trypsin-induced de-adhesion. (B) Schematic representation of axonal contraction concomitant with tip retraction upon de-adhesion. L_s_ is the straight line distance between two ends at time 0. L_0_, L_t_ and L_f_ are lengths in the first frame, an intermediate frame and the final frame, respectively. (C-C”) Representative frames from time-lapse imaging of an axon straightening following trypsin-induced deadhesion. While the growth cone and cell body do not show significant movement the axonal segment straightens. Trypsin is added at time 0. Time stamp shows minutes:seconds elapsed. Scale bar: 15 μm.

**Figure 2.**
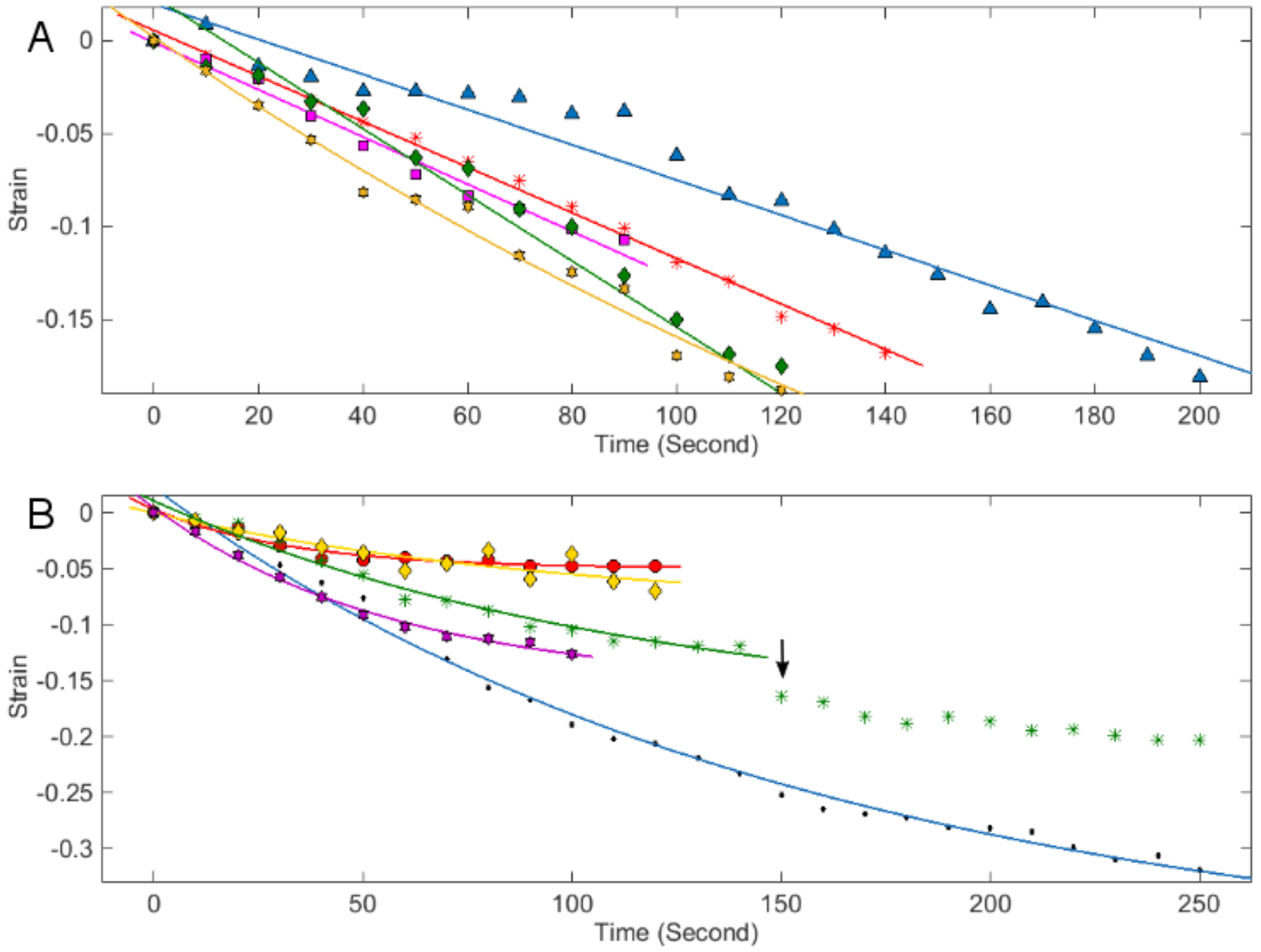
Axonal strain relaxation upon de-adhesion. Examples of strain relaxation of axons upon de-adhesion showing both (A) linear and (B) non-linear responses. Strain = (L_t_ − L_0_) / L_0_, where L_0_ and L_t_ are lengths at time 0 and at time t, respectively. Different axons are marked in different colors. Symbol indicate actual data while continuous lines are linear and single exponential fits. Arrow indicates sudden drop in the strain in that axon as an effect of delayed local detachment; hence data points further are not included in the fit.

### Axonal straightening after de-adhesion is actomyosin dependent

To investigate the contribution of actomyosin contractility in axonal straightening, a specific small molecule inhibitor of myosin II, Blebbistatin, was employed. Curved neurons pre-treated with 30 μM Blebbistatin for 1 hour failed to straighten upon trypsin-induced de-adhesion (Figure 3B-B””, C; n=10). Matched DMSO controls, on the other hand, showed the expected axonal straightening (Figure 3A-A””; n=9). To directly compare the two treatments, We use a time-independent parameter for the extent of contraction. Comparison of this contraction factor (see Materials and Methods for definition) between DMSO and Blebbistatin treated also indicated that inhibition of myosin II prevented deadhesion induced axonal straightening (Figure S2; p= 0.0001).

These experiments suggested that axonal strain relaxation observed upon axonal deadhesion is due to contractility of axons driven by myosin II activity.

We next evaluated the contribution of axonal F-actin by using Latrunculin A (Lat A), which depolymerizes F-actin by binding to actin monomers and preventing them from polymerizing. Lat A (0.6 μM) pretreatment for 15 mins was used to inhibit actin polymerization prior to trypsin-induced detachment and resulted in an expected decrease in phalloidin staining intensity along the axon (Figure S3 B). Lat A pretreatment completely abolished axonal contractility (Figure 4B-B””; n= 7) while matched DMSO controls showed straightening upon trypsinisation (Figure 4A-A””; n= 7). Direct comparison of contraction factors also indicated that axonal F-actin is essential for neurite contractility (Figure S3 A; p<0.0001).

We imaged the Blebbistatin and Lat A-treated neurons for up to 10 minutes to find out if axonal contractility was delayed. However, no late response was observed for both Blebbistatin and Lat A treated neurons. To ensure trypsin de-adhesion was not affected by drug treatment, fresh medium was flowed into the culture dish at the end of the imaging period to generate flow disturbances. Axonal segments were found to be detached and floppy indicating de-adhesion (data not shown).

Taken together these experiments implicate active actomyosin contractility in mediating axonal straightening. Actomyosin activity driven pulling forces have been previously demonstrated in axonal retraction (19, 27). Consistent with the significant actomyosin contribution observed in our studies, disruption of F-actin has been previously reported to reduce axonal rest tension (20). In fly motor neurons too axonal contraction is driven by actomyosin contractility (21) supporting our results using a different experimental paradigm in vertebrate neurons.

Unlike the above mentioned reports, our study avoids local/acute perturbations that may trigger axonal entry of extracellular calcium and in turn induce actomyosin contractility. The current study, using a strategy to probe axonal mechanics without localized maniulations, further underscores the importance of actomyosin activity in mediating axonal contractility.

**Figure 3.**
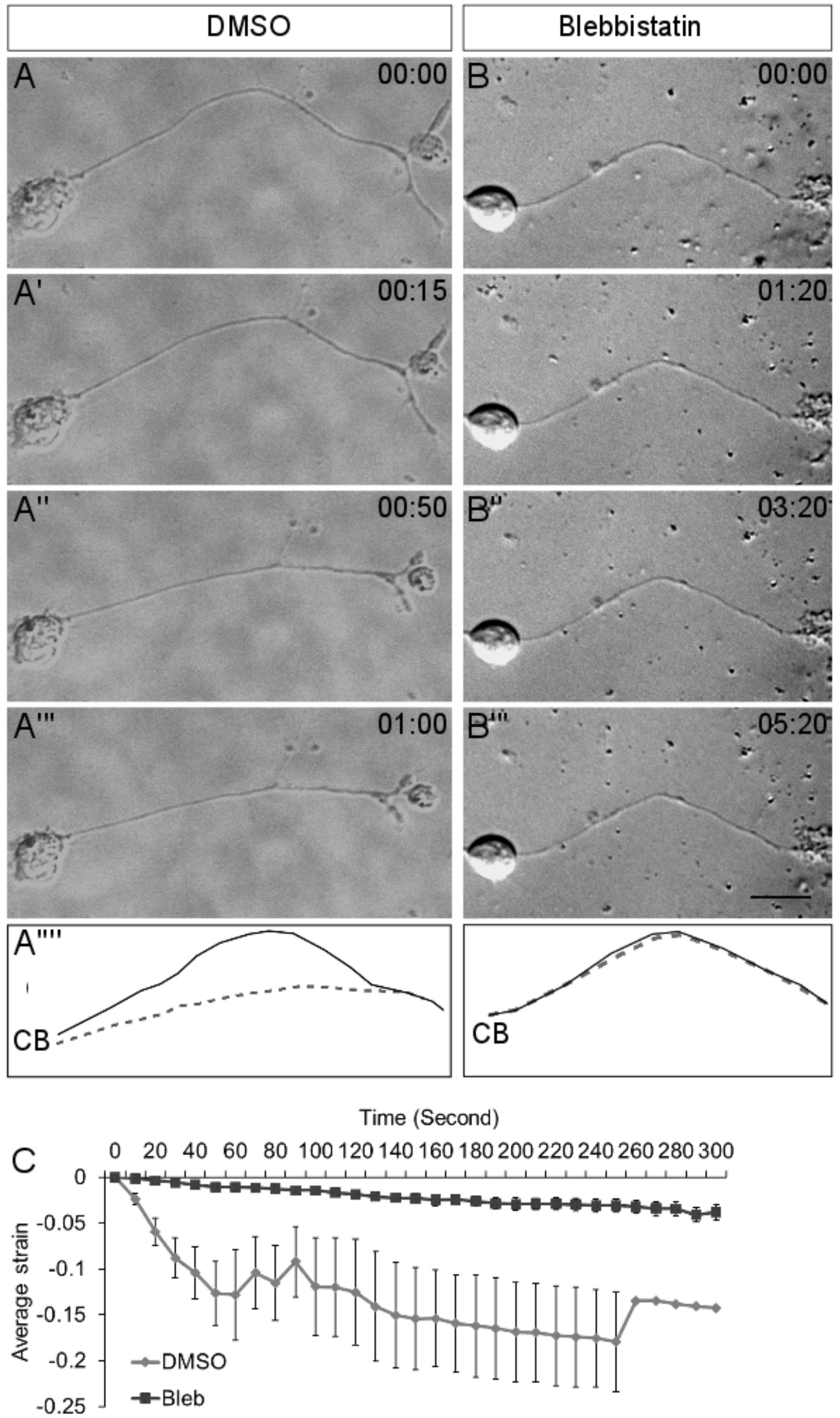
Axonal contraction is dependent on myosin II activity. (A-A”’) Representative frames from time-lapse imaging of a DMSO-treated control axon. (B-B”’) Representative frames from time-lapse imaging of an axon pretreated with Blebbistatin (30 μM) for 1 hour before trypsin addition. (A””), (B””) show the line trace of the axon shown in the first frame (solid line) and last frame (dashed line) of the representative micrographs of both treatments. CB indicates the position of the cell body. Trypsin is added at time 0 for each treatment. Time stamp shows minutes:seconds elapsed. Scale bar: 15 μm. (C) Average strain rate is reduced upon Blebbistatin (Bleb) treatment (n = 10) compared to DMSO-treated controls (n = 9). Error bars indicate standard error of the mean.

### Effect of microtubule depolymerization on axonal contraction

The axonal shaft is filled with a cross-linked bundle of microtubules, which contribute to multiple aspects of axonal behavior and mechanics (28). To test the role of the microtubule cytoskeleton in axonal contractility, neurons were pretreated with the microtubule depolymerizing agent, Nocodazole (Noco; 33.3 μM for 15 mins). Interestingly, Nocodazole pretreatment did not block axonal contraction (Figure 4C-C””; n=9) though the rate of contraction appeared to be reduced compared to the DMSO control (Figure 4D; n=7). Direct comparison of the contraction factor (see Supplementary methods for definition) also indicated that axonal contractility is affected upon Nocodazole treatment (Figure S3 A; p= 0.0204). Similar experiments using a lower dose of Nocodazole was consistent with the earlier results. Pre-treatment with 16 μM Nocodazole did not abolish axonal contraction (Figure S4 B-B”) as compared to matching DMSO controls (Figure S4 A-A”). The extent of contraction is comparable (Figure S5, D) however, the rate of contraction was reduced (Figure S4, C). Both doses of Nocodazole treatment resulted in a reduction in alpha-tubulin staining intensity (Figure S3C, S4E) as expected for microtubule disruption.

A recent study in *ex vivo* preparations of fly embryos, disruption of microtubules led to faster axonal contraction of motor neurons in response to slackening (21). In our study, microtubule depolymerization did not prevent axonal shortening but the rate of contraction was slightly reduced. This discrepancy may be due to intrinsic differences between invertebrate and vertebrate neurons. In addition, Nocodazole treatment resulted in most neurites exhibiting a beaded morphology (Figure 4C). This is consistent with previous observations (29) and confirms the efficacy of Nocodazole. This significant change in axonal morphology may adversely affect the contractility by indirectly affecting the continuity of the actomyosin network. Further, Nocodazole-treated axons also showed thin membrane tethers as they retracted (Figure S3 D-D’). Similar observations have been made previously (29) and it is likely that these tethers hinder the contraction of Nocodazole-treated neurons.

As axons self-shorten under actomyosin driven contraction, microtubules appear to be under compression and possibly accommodate the length reduction by sliding against each other (29). The slightly reduced contraction rate seen upon microtubule depolymerisation is probably a secondary effect of altered morphology (Figure S4) which precludes a clear evaluation of microtubule function in our experimental system.

It is possible that curved bundles of microtubules in curved axons play an active role in straightening in conjunction with the actomyosin machinery. However, whether individual microtubules exist in curved configurations in axons is unknown. Further, our experiments do not rule out the possibility that microtubule de-polymerization indirectly disrupts the axonal actomyosin organization and thereby damps the contractility. In the future, studies targeted at evaluating the contribution of mechanisms regulating microtubule sliding and force-dependent remodeling and subsequent reassembly to axonal contractility are likely to generate important insights.

**Figure 4.**
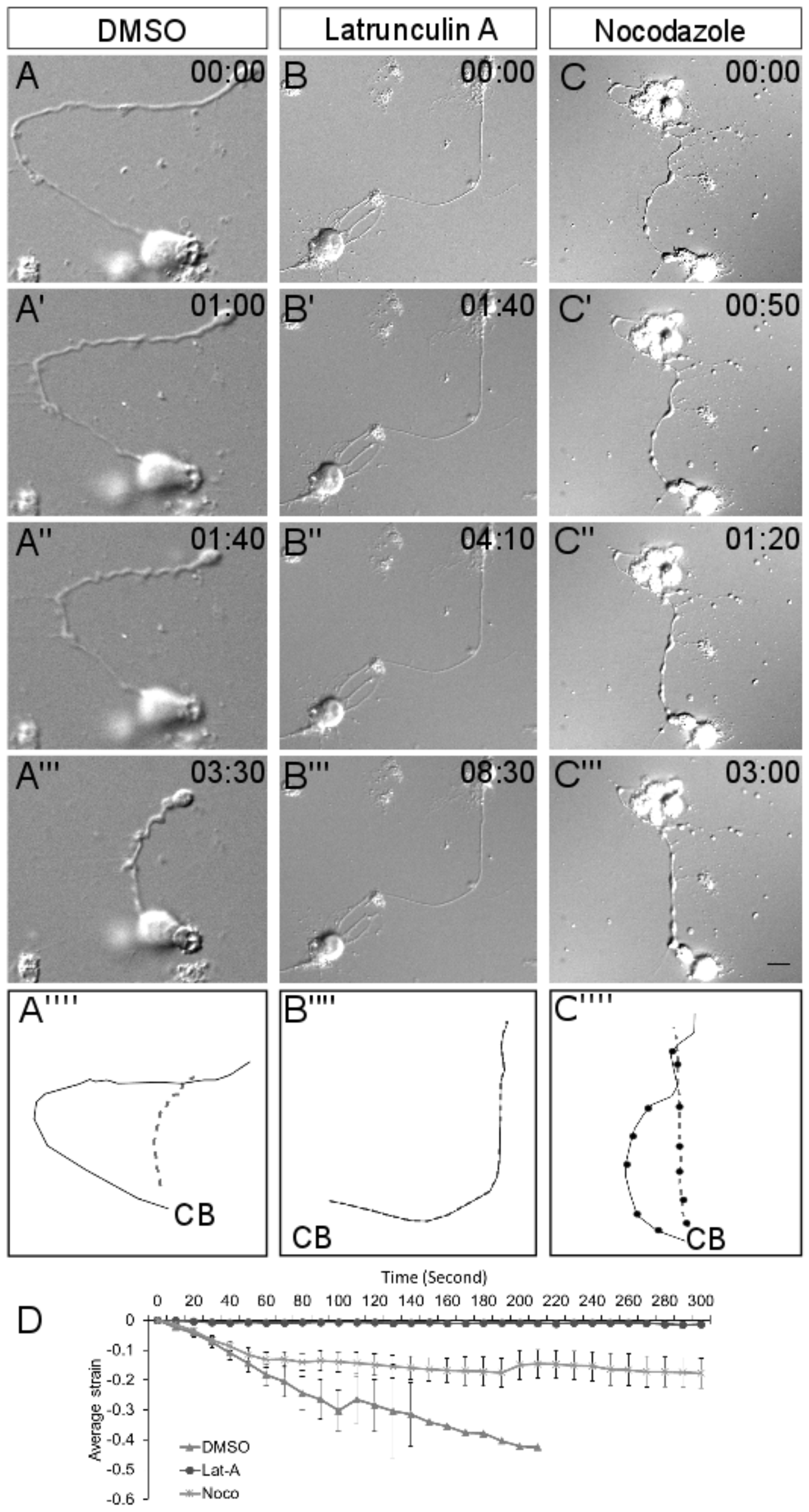
Effect of F-actin and microtubule depolymerisation on axonal contraction. (A-A”’) Representative frames from time-lapse imaging of a DMSO-treated control axon. (B-B”’) Representative frames from time-lapse imaging of an axon pretreated with Latrunculin A (0.6 μM) for 15 minutes before trypsin addition. (C-C”) Representative frames from time-lapse imaging of an axon pretreated with Nocodazole (33 μM) for 15 minutes before trypsin addition. (A””), (B””) show the line trace of the axon shown in the first frame (solid line) and last frame (dashed line) of the representative micrographs of the three treatments. CB indicates the position of the cell body. Trypsin is added at time 0 for each treatment. Time stamp shows minutes:seconds elapsed. Scale bar: 15 μm. (D) Compared to DMSO controls (n = 7), average strain rate is strongly reduced upon treatment with Latrunculin A (Lat-A; n = 7). Treatment with Nocodazole (Noco; n = 9) also reduces axonal contractility. Error bars indicate standard error of the mean.

### Cytoskeletal strain is heterogeneous along the axon

To evaluate cytoskeletal deformations upon trypsin-mediated de-adhesion and straightening, we tracked the position of labeled docked mitochondria present along axons. Mitochondrial interactions with actin, microtubules and neurofilaments have been reported previously and form the basis of several studies involving use of docked mitochondria as a marker for cytoskeletal deformations (30–32). In neurons, docked mitochondria have been established as a reliable marker for cytoskeletal dynamics in stretch (33) and elongation (34) paradigms. Mitochondrial responses are consistent with other readouts for cytoskeletal dynamics in these studies.

MitoTracker labeled axons were used in our straightening assay to investigate local cytoskeletal dynamics (Figure S5). Tracking of mitochondria using conventional kymography tools was not possible since the axon itself shortens. We used a custom written code to track position of mitochondria and calculate local strains between adjacent mitochondria pairs as a function of time. This local strain in this analysis is defined as: 
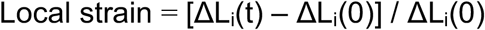

Where ΔL_i_ is distance between the i-th pair of adjacent mitochondria (indexed sequentially from one end) and t is time.

The variations in local strain did not show any observable distal to proximal trend (n=8). The local strains for pairs of mitochondria for an axon is shown in Figure 5 (inset shows the axonal strain). While the cumulative sum of the local strains was negative, at any given time both positive and negative strains were observed along the same axon and these values fluctuated with time (plot showing instantaneous strains for the same mitochondrial pairs is shown in Figure S7A). Interestingly, there were several instances where a mitochondrial pair was contracting but the other pairs were observed to be dilating (Figure 5, S7A). These data suggest that the axonal cytoskeletal in not contracting as an uniform whole but is highly heterogenous.

In order to distinguish the strain dynamics upon contraction from baseline mitochondrial fluctuations, we imaged mitochondria labelled axons without inducing trypsin-mediated de-adhesion (Figure S6A-A”). Minimal strain fluctuations at similar time scales were observed in these experiments (Figure S6B). As expected, average instantaneous strain was significantly different and negative in axons undergoing straightening as compared to axons not undergoing straightening (no trypsin-mediated de-adhesion; Figure S7B). Comparison of variances between axons undergoing straightening to those not treated with trypsin also revealed an increase in inter-mitochondrial strain variance confirming the heterogenous response of the cytoskeleton over and above baseline fluctuations in mitochondrial positions (Figure S7C).

Similar heterogeneous nature of cytoskeletal strain has been reported previously in axons under imposed uniaxial stretching (33). This study suggested that the location of axonal adhesion sites with the substrate could contribute to the heterogeneity in subcellular strain. However, in our assay, axonal segments are detached from the substrate and therefore preclude this possibility. Rather our study suggests that differences in local material properties of the cytoskeleton, including spatiotemporal variability in acto-myosin activity, may contribute to the strain heterogeneity observed during axonal contraction.

*In vivo*, the heterogeneous response seen along axons might be important for localized regulation of various processes like branch, adhesion dynamics and fasciculation. So far axonal contraction studies have been largely limited to the evaluation average properties like for force or bulk strain. We show that actomyosin-based contractility drives axonal contraction but the subcellular response is heterogenous.

**Figure 5.**
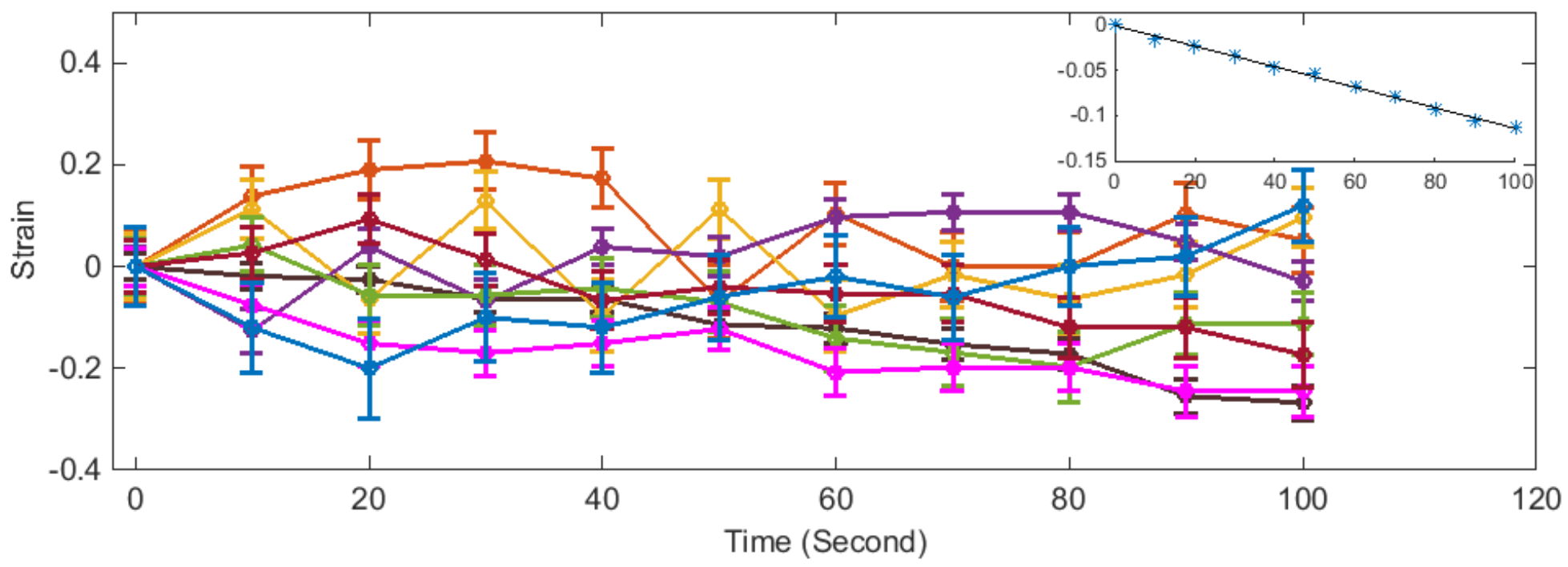
Intermitochondrial strains show heterogenous response. Plotting the evolution of strain between mitochondrial pairs (adjacent pairs numbered from soma to growth cone) of a randomly selected axon as it straightens. The data reveals instances of contraction in one pair of mitochondria being concomitant with an extension in another pair and occasionaly contraction/extension in two pairs occur together. For example, the purple and blue pairs, and also the orange and dark pink pairs. Inset shows that the net axonal strain decreases steadily with time compared to the local strains. Error bars are standard deviations of natural mitochondrial fluctuations observed in six live axons recorded for a similar period of time without inducing de-adhesion.

### Axons show spontaneous contractility between isolated adhesion sites

Axonal trajectories and configuration are determined by the directional translocation of the growth cone, that can be biased by local differences in environmental cues, and the resistance of the axonal shaft to bending (35). The intrinsic contractility of neurites would tend to result in straight axons unless this is offset by attachments with the substrate along the axon or branches. In trypsin-mediated detachment experiments, attachments along the axons are removed allowing the axonal actomyosin machinery driven contraction to minimize length and curvature. We wanted to investigate if axons growing *in vitro* also show spontaneous straightening due to intrinsic contractility without any external perturbation, like typsin mediated de-adhesion.

To investigate spontaneous axonal contractility, we grew neurons on patterned substrates with isolated islands of extracellular matrix proteins (laminin and fibronectin).These experiments were done in serum containing media. These islands represented isolated sites of high-adhesion relative to the intervening space. We observed that axonal segments often had initially curved trajectories between two islands. However, with time, axonal contraction was established in the segment spanning the two islands resulting in the reduction of curvature and length (Figure 6; n=8; Movie S2). In examples where one end of the axonal segment was the growth cone, axonal contraction resulted in straightening of the segment without any obvious growth cone movement (Figure 6A-A”). This precludes growth cone towing driven tension buildup on the axon and suggests that axons are intrinsically contractile. We observed straight axonal segments for substantial periods of time and confirmed that straight axons remain straight and the observed spontaneous straightening are not random shape changes (data not shown).

The spontaneous shortening observed in these experiments progressed slowly, typically taking tens of minutes-to an hour or two, which is significantly slower than trypsinization-induced contraction (Figure 6B). This delay suggests that detachment along axons is rate limiting in unperturbed axonal contractions. As the contraction factor (see Materials and Methods for definition) is a parameter for the extent of contraction, we used it to compare between spontaneous and de-adhesion-induced contractility. The contraction factor was comparable between the two groups (Figure 6C) suggesting a similar degree of contraction in the two assays.

Straightening of axonal segments without appreciable growth cone movement implicates axon intrinsic contractile mechanisms, as opposed to growth cone generated forces in axonal length minimization. A previous study, using locust neurons, has described a similar assay using islands of carbon nanotubes on quartz sheets. Axonal segments were found to straighten between islands and branches along the contracting segment were retracted concomitant with the development of tension (8).

**Figure 6.**
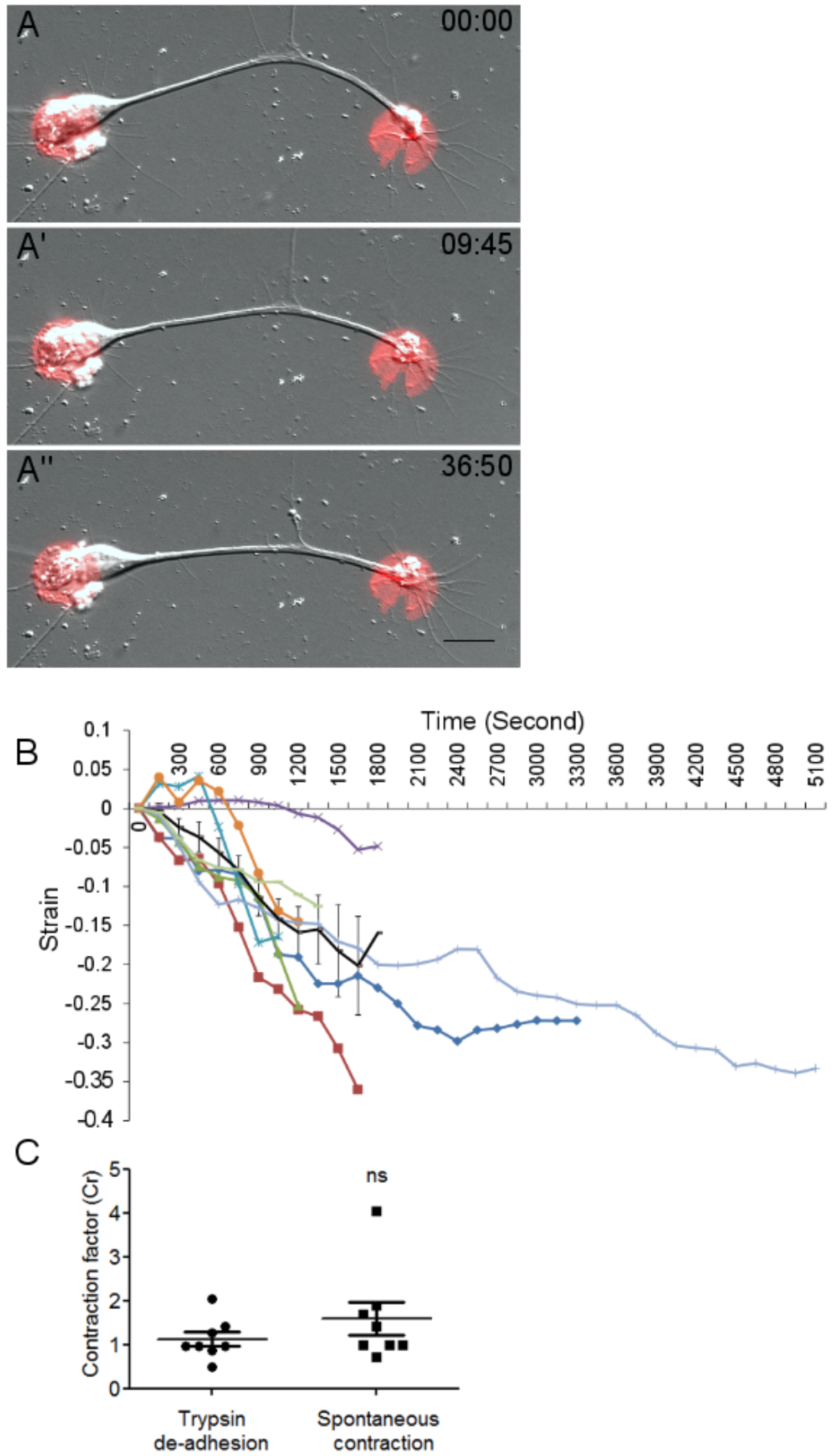
Spontaneous contraction of axons. (A-A”) Representative micrographs from time-lapse imaging of a DRG axon spontaneously straightening between two adhesive islands containing laminin and fibronectin (red circles). Time stamp shows minutes:seconds elapsed. Scale bar: 15 μm. (B) Evolution of strain with time for 8 axons and the average strain plotted with error bars indicating standard error of the mean. In some axons, positive strain is initially observed before the axon begins to shorten (Figure 6B). This is because this assay relies on slow, spontaneous de-adhesion unlike acute induction of de-adhesion by trypsin. (C) Comparison of Contraction factor (see Materials & Methods for definition) suggest that the extent of spontaneous contraction between adhesive sites (n = 8) is comparable to that observed during trypsin-induced de-adhesion (n = 8). Unpaired t-test was used to compare the means. Error bar represent standard error of the mean. ns: not significant.

## CONCLUSIONS

We have developed a simple assay to evaluate axonal contractility that avoids local, acute mechanical perturbations and instead uses trypsin to detach curved axonal segments from the substrate. In addition, we have developed a microcontact printing based assay to assess spontaneous tension development along axonal segments.

Collectively, our data suggest that axons are inherently contractile and tend to minimize their length unless opposed by adhesions along their length. Upon release of adhesions, intrinsic contractility drives the system towards minimal length. Our study implicates the actomyosin system as the major driver of contractility. Analysis of subcellular strain using docked mitochondria to evaluate local axonal strain shows that the contractility is heterogenous along the axonal length.

This study highlights the necessity to identify the detailed organisation of the the axonal acto-myosin system. Recent advances in super-resolution microscopy has revealed a membrane-associated periodic skeleton (MPS) comprising of alternating ßII spectrin and F-actin rings (12, 13). Using STED nanoscopy we evaluated the prevalence of ßII spectrin (immunostained with an anti-ßII spectrin antibody) and F-actin (live axons stained with the cell permeable actin probe, SiR-Actin) MPS in chick DRG axons grown for 2 days *in vitro* (DIV). These studies revealed that ßII spectrin (Figure S8) and F-actin (Figure S9) MPS were not prevalent in 2 DIV chick DRG axons with only a weak periodic organisation (Figure S8, S9). In contrast, later stage axon (5 DIV) had clear and extended MPS along the entire axon length (Figure S8E, S9E). Our observations are in line with observations in mammalian neurons in culture where MPS is abundant only in late stages (DIV 10) while earlier stages do not show such organisation (36, 37). At 2 DIV, F-actin labelling is diffuse and inhomogeneous along the axon. F-actin patches and elongated bundles were occasionally observed. We further evaluated myosin distribution using an antibody against phosphorylated myosin light chain (pMLC) and STED nanoscopy. pMLC was found to have a punctate distribution along the length of the axons (Figure S10). A recent study had also reported pMLC distribution along the entire length of 3 DIV rat hippocampal axons (38). The latter study and our results also indicate that pMLC is not commonly colocalised with the F-actin patches.

Spectrin is known to be sensitive to axial tension and has been demonstrated to regulate rest tension in worm neurons (14, 39) Our observation that MPS is not abundant in 2 DIV axons does not rule out the function of a non-periodic membrane-associated spectrin-actin network. Both ßII spectrin and actin are distributed along the axonal cortex and are likely to influence the mechanical properties of axons even in early stages. The prominent organisation of actin in periodic rings in older neurites raises the question how such structures may support axial contractility. This question will have to be systematically investigated in the future, especially in light of recent results showing that in older neurons pMLC co-localizes with actin rings in the axon initial segment while being conspicuously absent from the distal axon (38).

In summary, while these data do not identify specific acto-myosin organisation contributing to axial contractility, they suggest that in 2 DIV chick DRG a diffuse network of cortical acto-myosin organisation may regulate contractility along the length of the axon (Figure 7).

As the axonal length shortens, it is expected that microtubule sliding will be necessary to accommodate this change (Figure 7). Thus, future investigations into the function of microtubule sliding and crosslinking activities are also likely to reveal important mechanistic modalities.

In a complex environment, turning and pausing of growth cone might impart curvature along axons. Branch dynamics and competition between the branches can also lead to curved morphology of axons. During neurite outgrowth the growth cone translocation results in pulling forces that is balanced by the neurite tension. It has been suggested that modulation of this force balance may influence growth cone turing (40). In addition to growth cone mediated straightening (41), passive stretch from surrounding tissue can also contribute to axonal tension (42). In our experiments, straightening on patterned substrates suggests that while axons may follow curved trajectories due directional biases of the growth cone, the axonal contractile forces are able to spontaneously develop and straighten axonal segments between two points of strong adhesion. This occurs without towing by the growth cone and points to axon-intrinsic contractility as major regulator of axonal tension and in turn axon conformation. Though currently unknown, it is possible that the adhesive sites may initiate mechanochemical feedback responses that enhance axonal contractility. We propose that spontaneous acto-myosin driven axonal contractility as a local mechanism for maintaining the rest tension, which is known to influence various neurodevelopmental processes, including length minimization at the level of network.

**Figure 7.**
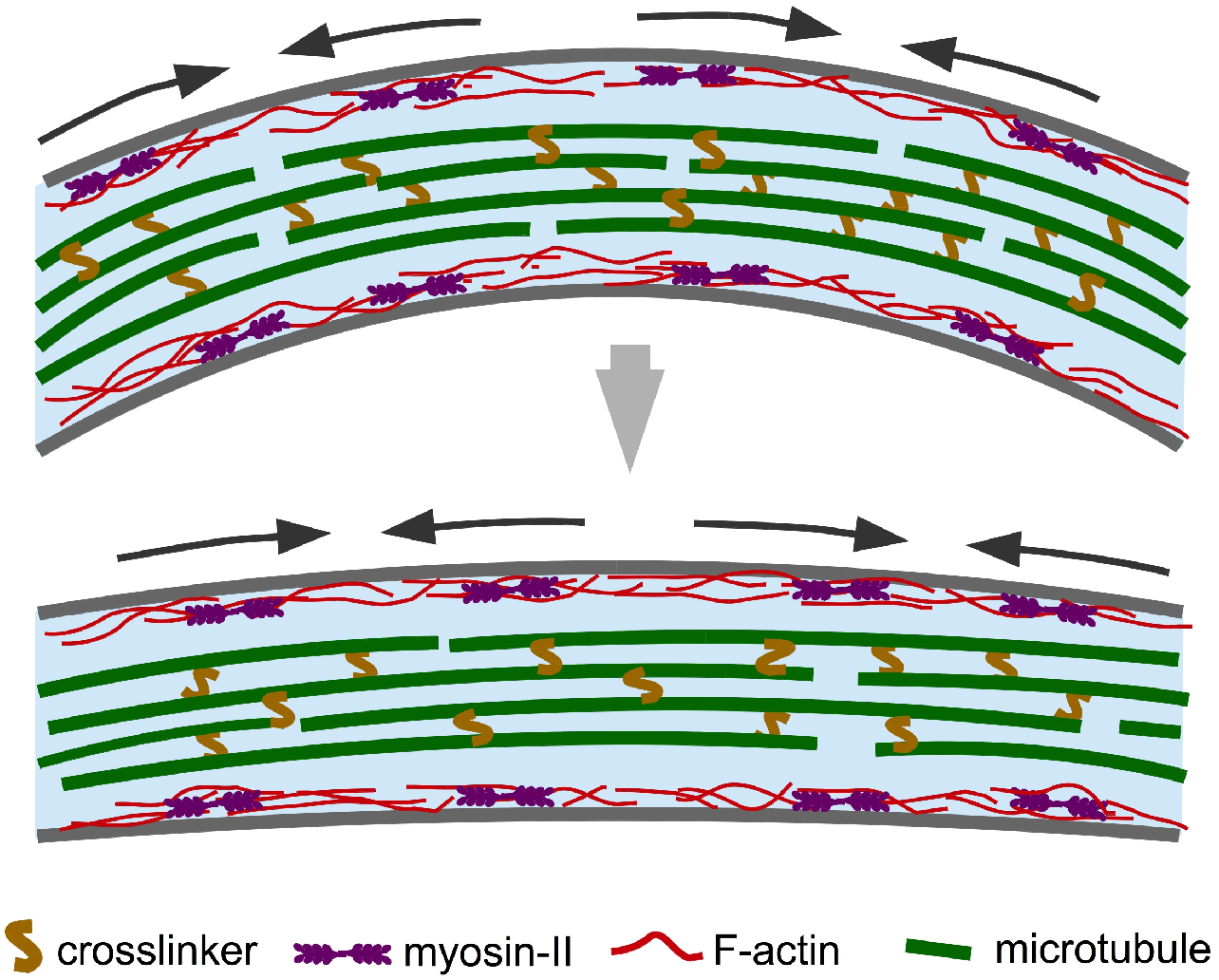
Model for cytoskeletal mechanisms mediating axonal straightening. In 2 DIV chick DRG neurons, the cortical actomyosin network contracts upon detachment induced by unknown feedback mechanism leading to the straightening of axons. The acto-myosin network appears to be distributed along the length of the axon, capable of local remodelling (indicated by pairs of arrows). A weak and discontinuous periodic actin-spectrin skeleton at this stage and is not shown in this schematic. Microtubule sliding and rearrangement are likely to be necessary in order to accommodate the contraction.

## AUTHOR CONTRIBUTIONS

S.M., P.P., J.J. and A.G. designed the research and analysis. S.M. performed the research and analyzed the data. S.M., P.P., J.J. and A.G. wrote the manuscript. A.G. secured funding. All authors read and reviewed the manuscript.

## ACKNOWLEDGEMENTS

The authors acknowledge intramural funding from IISER Pune and an infrastructural Nano Mission Council, Department of Science and Technology [SR/NM/NS-42/2009] grants to IISER Pune. The IISER Pune Microscopy facility is gratefully acknowledged for access to microscopes. Dr. Richa Rikhy, IISER Pune, is acknowledged for providing MitoTracker Green FM. Dr. Deepak Barua, IISER Pune, is acknowledged for advice regarding statistical analysis. The authors thank Prof. N. K. Subhedar for critical reading of the manuscript.

